# Deep Learning assisted Peak Curation for large scale LC-MS Metabolomics

**DOI:** 10.1101/2020.08.09.242727

**Authors:** Yoann Gloaguen, Jennifer Kirwan, Dieter Beule

## Abstract

Available automated methods for peak detection in untargeted metabolomics suffer from poor precision. We present NeatMS which uses machine learning to replace peak curation by human experts. We show how to integrate our open source module into different LC-MS analysis workflows and quantify its performance. NeatMS is designed to be suitable for large scale studies and improves the robustness of the final peak list.

## MAIN

Liquid chromatography-mass spectrometry (LC-MS) is a widely used method in untargeted metabolomics. The post-acquisition raw data processing which aims to detect compound related peaks and distinguish them from noise signals is still a major challenge. Many algorithms and tools have been developed to address this challenge (e.g. XCMS^1^, MZmine^2^, Optimus^3^). Pipelines for the automatic LC-MS raw data processing usually consist of the following steps: definition of regions of interest (ROI), detection of chromatographic peaks, quantification of these peaks, peak matching or grouping for samples within the batch or analysis, clustering of peaks belonging to the same compound. XCMS and MZmine are the most widely used open source software which perform all these steps and provide the user with a table of peaks found in the spectra and their integral intensities for each sample. However, there is a tendency for peak picking software to over-pick peaks (i.e. creating a high number of false positives^4^), poor consistency between software is another major issue^5^. Both issues may obstruct or impede downstream analysis and biomedical interpretation of metabolomics studies and thus some kind of manual peak curation is still the norm. This also makes analysis of large scale studies extremely laborious and limits reproducibility of analysis. Recent progress in machine learning (ML) algorithms^6^ and availability of affordable parallel processing hardware (GPUs) has sparked application of deep learning methods in both GC-MS^7,8^ and LC-MS in peak detection^9,10^. ML has also been used for intra and inter batch correction^11^ in LC-MS.

Here we introduce NeatMS which is designed to serve as an independent deep learning based peak filter tool in existing analysis pipelines. It addresses the over-picking issue by automatically evaluating and classifying peaks based on quality. To achieve this, we introduce three peak quality classes (high, acceptable, “poor quality or noise” – henceforth called noise) and provide a pre-trained neural network to allow for out of the box usage. Transfer learning and complete retraining are also supported, see methods. NeatMS can be easily integrated into existing workflows, see Figure 1.a, it uses a convolutional network architecture shown in (Figure 1.b). Further algorithm details are given in the methods section.

**Figure 1:**
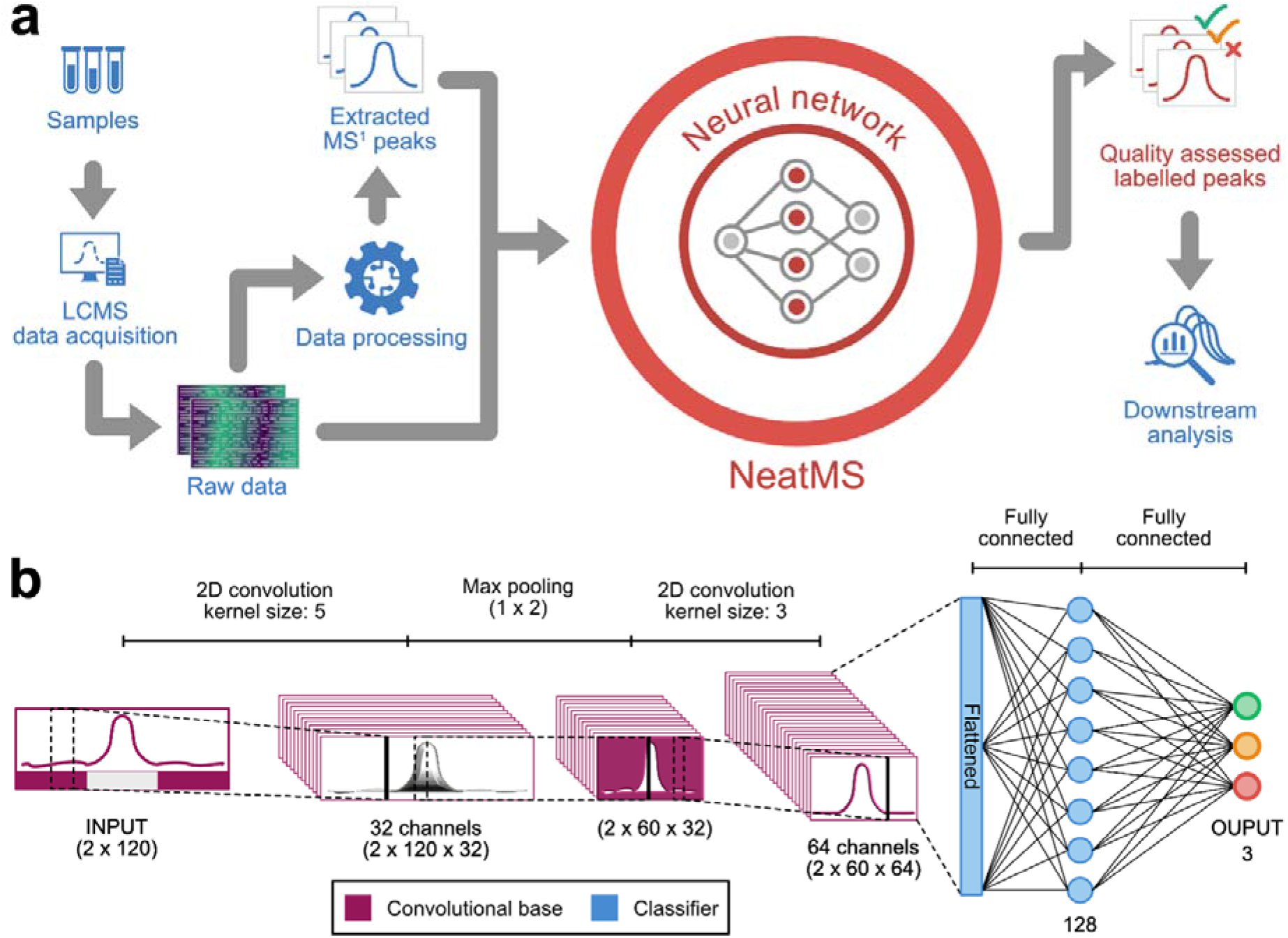
**(a)** Integration of NeatMS (red) into an existing workflow (blue): Samples are run through the LCMS data acquisition, raw data is processed using standard automated tools to extract peaks, both raw data and extracted peaks are used as input for NeatMS that assigns them to three classes: high quality, acceptable or noise. The classification information can be used in the downstream analysis. NeatMS comes with a pre-trained network and includes all components for retraining and transfer learning. **(b)** Architecture of the neural network: NeatMS includes a 2D convolutional base for feature extraction and a classifier made of two fully connected layers. This architecture was chosen due to its high performance in object detection and pattern recognition^12^. The max pooling layer between the convolutional layers reduces data size in the retention time dimension. This enables a higher abstraction of the data and reduces the number of learned parameters and thus helps to prevent overfitting as well as to reduce computational training effort. The classifier is made of two fully connected layers and uses a softmax function to produce three output values which correspond to the peak classes.

To evaluate the performance of NeatMS we use two data sets with known chemical standards (CS). Dataset 1 consists of 20 quality control samples from the Biocrates P400 kit^13^ consisting of 80 chemical standards (CS) at proprietary concentrations in a lyophilized plasma matrix. Dataset 2 is based on the Biocrates kit calibration sample “Cal 1”, which is matrix free and contains 41 chemical standards. We created a serial dilution in water (1:1.2, 1:1.4, 1:1.6, 1:1.8, 1:2, 1:3, 1:5, 1:7.5, 1:10, 1:15, 1:20, 1:30, 1:50 and 1:100). Following dilution, we added 39 compounds (Biocrates internal standard mix) at the same concentration to each sample to act as internal standards. Each dilution was measured in triplicate. The object of this dilution series was to objectively assess how NeatMS performed over a range of peak intensities. All data was acquired following Biocrates P400 kit standard protocol on an Agilent 1290 coupled with a Thermo QExactive instrument.

Our NeatMS analysis first evaluates the pre-trained (PT) model. This initial model was trained on a wide range of peak shapes (see methods). Additionally, we used transfer learning to adapt the PT model to the actual dataset 1 by manually assigning a small subset of peaks to the correct quality class and using an 80/10/10 training/test/validation approach (see methods for details); we will call the second model TL. The PT as well as the TL model are provided as supplementary material. All components needed to perform model training and transfer learning are part of the open source NeatMS software. We processed MZmine and XCMS centWave^14^ peak-picking output for set 1 and 2 using NeatMS, employing both the PT and the TL models. Table 1 summarizes the number of peaks and class assignments for dataset 1. Furthermore, the table shows results for the recently published peakonly tool, which also applies machine learning on raw data to detect high quality peaks. Table S1 summarizes all parameters and software versions used and the methods section describes how they were chosen. Further discussion will be focused on NeatMS using the MZmine trained TL model. Results with XCMS could likely be further improved by creating an XCMS-specific trained model.

**Table 1:** Tool and model comparison using dataset 1 showing average number of peaks found across 20 samples and average percentages of detected CS. The input row shows the results returned by the original peak detection tools (MZmine, XCMS), other rows show the details of the three peak classes given by NeatMS. The total number of peaks after classification is smaller than the input due to the application of a minimum scan number filter that NeatMS uses (default value of 5 is used).

|  |  | NeatMS<br>MZmine TL<br>model | NeatMS<br>MZmine PT<br>model | NeatMS<br>XCMS TL<br>model | NeatMS<br>XCMS PT<br>model | peakonly |
| --- | --- | --- | --- | --- | --- | --- |
| Peak<br>number | Input | 6977 | 6977 | 5994 | 5994 | 1907 |
|  | Classified | 5505 | 5505 | 4513 | 4513 | Not<br>reported |
|  | High Quality | 1069 | 2127 | 2280 | 2635 | Not<br>reported |
|  | Acceptable<br>Quality | 1945 | 1817 | 714 | 1088 | Not<br>reported |
|  | Noise | 2491 | 1560 | 1519 | 791 | Not<br>reported |
| CS found | Input &<br>classified | 94.25% | 94.25% | 97.13% | 97.13% | 79.44% |
|  | High Quality | 79.31% | 88.19% | 91.44% | 92.94% | Not<br>reported |
|  | Acceptable<br>Quality | 11.25% | 4.94% | 2.13% | 3.69% | Not<br>reported |
|  | Noise | (3.69%) | (1.13%) | (3.56%) | (0.50%) | Not<br>reported |

Median relative standard deviations (RSDs) increase substantially from high quality to noise peaks (Figure 2.b), with acceptable quality peaks falling in between. Our quality class assignment differs strongly from conventional QC-RSD filtering methods because we also observe many noise peaks with low RSDs. This effect is present with XCMS and MZmine for both models (not shown). Figure 2.d and 2.e show that the CSs are consistently found by NeatMS across various low dilution samples and generally tend to move from high to acceptable quality as dilution increases. Eventually some fall into the NeatMS noise category while most can no longer be detected by MZmine as the signal decreases with increasing dilution. Figure S1 (peak width distribution) shows that the noise class is dominated by rather broad peaks while the high quality class shows a consistent peak width distribution independent of the peak area. This indicates that our three classes represent a sensible quality classification for peaks. Figure 2.a (ROC curve) shows that the learning itself was very successful, the model closely resembles the expert knowledge of the trainer. Thus NeatMS evaluation is comparable to but much faster and much more reproducible and consistent than human expert evaluation. While an expert may still perform best for small numbers of peaks, NeatMS must be considered superior for large scale studies with hundreds or thousands of samples and potentially several millions of peaks. By including training and transfer learning functionality into our solution, we empower researchers to adapt the learned classification and filtering optimally to their specific data, needs and preferences.

**Figure 2:**
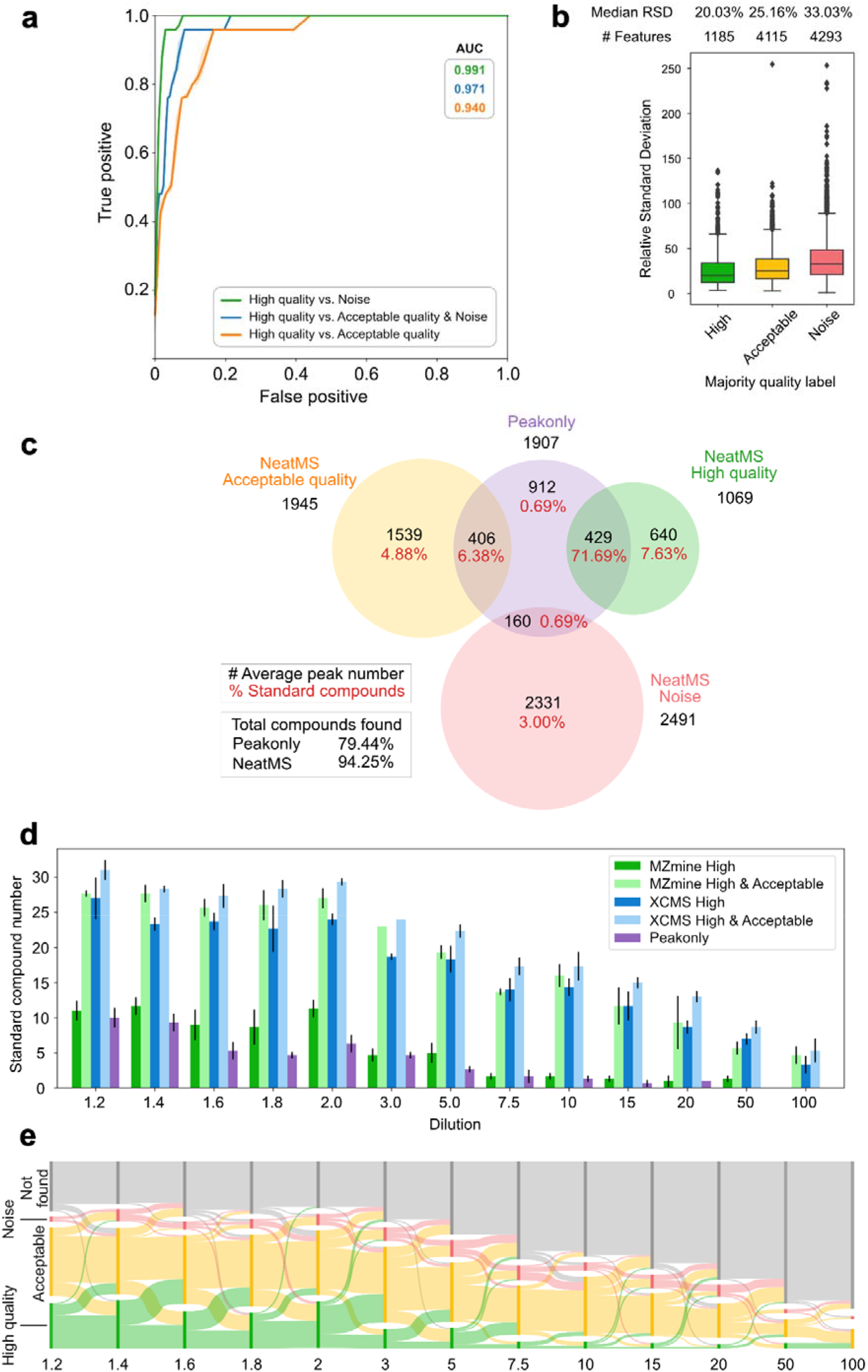
NeatMS results for dataset 1 (A, B, C) and dataset 2 (D and E) based on TL model and MZmine output. (**a**) ROC curves showing learning efficacy for three different group separations. The curves are created using the validation set of the TL model learning, high quality class probability returned by the model is used as the varying parameter. (**b**) Box plots of RSDs: High quality peaks show lower RSD and less RSD variability. Each feature for which peaks were present in at least 4 samples was assigned to a quality based on the most frequently reported class. (**c**) Venn diagram comparing peakonly and NeatMS. Numbers show averages over the 20 samples: total number of detected peaks (black), percent of recovered CSs (red). On average NeatMS reports 63.5 CSs as high quality, i.e. recovers 79% of original, another 11% are considered acceptable quality. Please note that matching peaks derived by different algorithms is challenging in itself and not completely unambiguous, cf. methods for details. (**d**) Classification of individual CS over different dilutions and for different tools, NeatMS high quality class outperforms peakonly for most dilutions. Acceptable quality class recovers a substantial number of additional CSs. (**e**) Sankey diagram showing the distribution of the 41 diluted CSs for all 3 replicates in different quality classes and their change between dilution steps. Each dilution step is represented by a stacked barplot, the widths of the flows between bar plots (dilutions) represent the fraction of CSs going from one class to another.

Table 1 shows that even if we do not use transfer learning NeatMS can still deliver a useful improvement for MZmine workflows. It substantially reduces the number of peaks that need to be considered for downstream analysis. This facilitates e.g. differential analysis either on only high quality or on high and acceptable quality data. For XCMS the separation between acceptable and high quality class is not so clean due to the abovementioned training approach. Both MZmine and XCMS users can start working with the PT model and will immediately improve their workflow performance and incrementally create further improvements by training better models.

Figure 2.d shows that NeatMS high quality class equals or outperforms peakonly in CS peak recognition for all dilutions. Additional CSs are classified as acceptable. A more detailed comparison for dataset 1 is shown in Figure 2.c (for XCMS we find comparable results, see Figure S2). NeatMS High and Acceptable quality classes together contain on average more than 90% of CSs. Peakonly reports an average of 1907 peaks, containing on average 79% of the CSs. Approximately the same percentage of CS is contained in 1069 high quality peaks reported by NeatMS. The concordance is however not perfect. Only rarely do we miss CSs that peakonly reports (0.7%). A small portion of the CS matched signals are considered Noise by NeatMS, upon visual inspection our expert usually agrees with the NeatMS algorithm (see Figure S5).

NeatMS requires neither high compute power nor long compute time; all data analysis described in this manuscript can be done with a standard laptop within minutes, all described training can be done with a modern PC within a few hours. NeatMS software is available as open source on github under permissive MIT licence and also provided as easy to install pipy and bioconda^15^ packages. NeatMS comes with a comprehensive user documentation, tutorials, and importantly also contains an easy to use training tool. Users can thus create their own models or improve existing one according to their specific needs. NeatMS supports standard input and output formats and is therefore easily added into existing workflows. Thus, it is compatible with many use cases and may help to enable improved and reproducible data analysis for large LC-MS studies.

## METHODS

Our software is designed to be easily integrated into existing workflows and is adaptable to different measurement protocols, instruments and preprocessing tools. Therefore, NeatMS can used in different ways. Most frequently it is used to classify an input dataset using an existing model. NeatMS also allows the creation of new or improvement of existing models using training data. The training data can be generated with an integrated labeling tool. For all usage the input data formats are the same and processing always starts with a data preparation step.

### Data Preparation

Although the peak detection is performed by an external tool, workflow or pipeline, the signal used for classification is directly retrieved from the raw data. This prevents biases originating from data transformations applied by the different peak detection tools (baseline correction, smoothing, etc…). Therefore, the input data of the module consists of .csv formatted files describing the peaks detected and the raw sample files in mzML format. Other vendor specific raw file formats can be converted into mzML format using the msconvert tool available in ProteoWizard^16^. The csv input files can be generated using standard preprocessing tools such as MZmine or XCMS. The position of the module within standard data processing workflows is illustrated in Figure 1.a. The output of NeatMS is again in csv format and contains the information from the input csv as well as the peak classification and labeling generated by the software.

Before any transformation NeatMS excludes unacceptably narrow peaks from further processing by requiring a minimum scan number of 5 (configurable minimum scan number input filter). As the model evaluates the peak shapes and the quality of their extraction from the raw signal (e.g. peak boundaries), it is important to provide contextual information. This is performed by extracting a larger retention time (RT) window to conserve the signal surrounding the peak (called peak margin thereafter). The RT window of the signal to be extracted is defined as follow (with *n* = 1 by default):

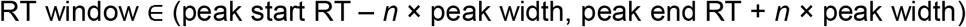

A min-max normalization is then applied to the extracted signal:

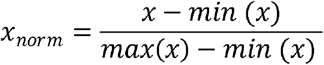

The resulting signal is linearly interpolated to obtain a vector of size *s* (120 by default). The s/(2n+1) that is, by default, the central 40 values, represents the peak signal and the n × s/(2n+1) that is by default the 40 values on each side represents the peak margins as shown in Figure 3. A second (binary) vector of length *s* is then created to describe whether an intensity value (single point) is part of the peak window (1) or the margin (0). The resulting data structure is a 2-dimensional tensor, or matrix, of size 2 × *s* as shown in Figure 3. Although *n* and *s* can be adjusted by the user, the pre-trained model provided with NeatMS has been trained using default parameters and require no adjustment when using this model.

**Figure 3:**
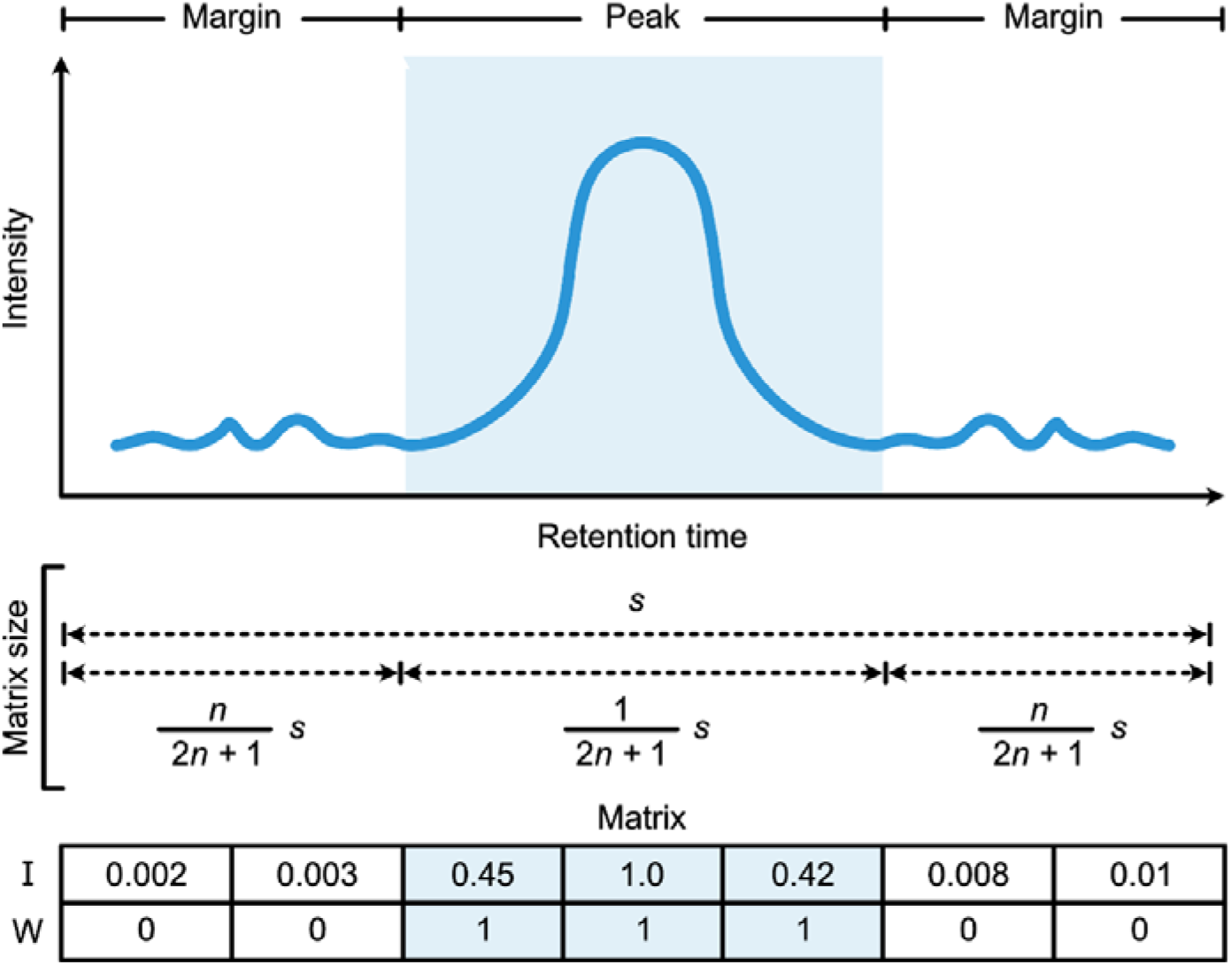
Data structure of a single peak after preprocessing for neural network feeding. ***s***: Size of scaled intensity vector. ***n***: Margin width (as a fraction of peak width). ***I***: Scaled intensity vector of size ***s***. ***W***: Binary window vector of size ***s*** (0=margin, 1=Peak).

### Convoluted neural network architecture

The neural network (see Figure 1.b) is built following generic convolutional network architecture including a convolutional base for feature extraction and a classifier made of fully connected layers. The convolutional architecture of the network was selected due to its high performance in object detection and pattern recognition^12^. The convolutional base is composed of two convolutional layers, with a max pooling layer between them. This operation reduces by half the size of the data in the retention time dimension and enables a higher abstraction of the data to classify, which helps to prevent potential over-fitting. This layer also reduces the number of parameters to learn and the computational cost. The classifier part of the network is made of two dense layers and produces three output values through a softmax activation function, which corresponds to the number of peak quality classes. ReLU (Rectified Linear Unit) activation function is used throughout the rest of the network. The convolutional layers use a stride of 1 with a “same” padding. Kernel size and channel number for every layer is detailed in the Figure.

### Training

As mentioned before NeatMS includes a pre trained base model. This PT model has been generated using a wide variety of peaks and can be used “as is”. However, it is possible to achieve models that are better suited to individual user needs by training the neural network on data generated by the specific column/instrument/peak detection workflow combination.

The creation of a new training dataset is facilitated by an interactive visualization and labeling tool that can be run within a Jupyter notebook^17^. This tool requires the same input as the main NeatMS tool and presents the user with randomly selected peaks for manual assignment to the three labeling classes. Typically, a few hundred peaks can be labeled within an hour. The PT model is based on about 5000 peaks, the TL model was adjusted using about 2500 peaks. Once the desired number of peaks have been labelled, the model can be trained using two different approaches (full training or transfer learning, see below). The labeled dataset is divided in an 80/10/10 training/test/validation split by default. Model testing and validation are always performed the same way regardless of the training approach chosen. The test set is used during the actual training process to prevent overfitting. The validation set remains untouched during the entire training process and is subsequently used for hyperparameter optimization. This optimization can be performed automatically or manually, but performing it manually allows more control over the specificity vs. sensitivity of the model. Instructions for manual process are given in the documentation.

The training tool enables the freezing of any network layer, making it possible to select the specific layers in which weights should be adjusted. It is, however, considered better practice to only adjust the classifier part by freezing the entire convolutional base when the training set is very small. Guidance on layer selection is given in the advanced section of the documentation. This approach is especially important for transfer learning and enables fine tuning of the pretrained models by further training specific layers^18^. The advantage of this approach over full training is that it requires a much smaller training dataset and thus less manual labeling effort. However, the tool also supports full training of entirely new models. This approach consists of using only the network architecture and fully training the model from scratch. This approach will produce the best results for the data being analyzed but requires a large training dataset. Instructions on how to import models are available in the documentation.

### Data Analysis Details

To evaluate NeatMS, dataset 1 and 2 were preprocessed with MZmine and XCMS using the versions and parameters detailed in Table S1. We choose the peakonly parameters as suggested by the authors and also tried to further optimize them, see Figure S4. Unfortunately, it was not possible to apply transfer learning or any retraining for peakonly because the software does not provide the necessary components to do this. For XCMS centWave we applied IPO^24^ for parameter optimisation individually for each dataset, furthermore we used the XCMS peak merging feature. For MZmine we used the parameters recommended in the user documentation.

Our validation with dataset 1 focuses on the number of known spiked chemical standards found by the different tools. Peak identification was performed using compound specific RT and *m/z* tolerance windows provided with the Biocrates kit. To compute the final results, percentages were calculated on the basis that all 80 compounds are detectable in all 20 samples. The same approach was used to analyse the recovery of CSs in dataset 2. These samples contain the same 80 CSs as dataset 1, but only 41 of the CSs were diluted while 39 were used as internal standards and thus were at the same concentration throughout (Figure S3.b). The Sankey diagram (Figure 2.e) was created by comparing the peak classes of the CSs for all three replicates in the consecutive dilution points to generate migration flows. A class corresponding to non-detected CSs was added to conserve an even CS population size throughout. To create the Venn diagrams (Figure 2.c and S2), peaks reported by one tool were compared to the list of peaks reported by the other tool and considered the same when any two peaks presented a mutual overlap higher than 50% in the RT and *m/z* dimensions, respectively. However, tools can differ widely in the peak boundaries assignment for the same peak. Therefore, any peak matching method will remain imperfect and ambiguous. This explains the non-complete overlap of peaks found by NeatMS when compared to peakonly. To generate RSD results in Figure 2.b we used peak alignment (using the MZmine join aligner algorithm) and assigned the features to the most frequently found quality class across the 20 samples. Features were retained only when present in a minimum of 4 samples.

### Module

NeatMS is written in python 3.6 and is available as a python package through pypi package installer and bioconda. The data handling and operations are performed using NumPy^19^ and scikit-learn^20^ and the neural network is constructed using Keras^21^ and Tensorflow^22^. As a python package, the intended use of the module is to be embedded as an extra step within a data processing pipeline. The module can be integrated and automatically executed by any pipeline or workflow management tool capable of running python code. However, it can also be used as a standalone application through a dedicated python script or within a Jupyter notebook. Several Jupyter notebooks are provided for tutorial purposes and can serve as templates and examples. The generated results are reported in standard .csv format and can also be exported as pandas^23^ dataframes for direct integration in python supported pipelines. The structure of the output can be controlled through a dedicated method to ensure smooth integration into the majority of data processing pipelines. Optional filters can also be turned on and parameterized. Details about full usage of the export method are provided in the documentation.

## Data availability

The github repository contains some sample data from dataset 1, the full datasets can be downloaded from http://doi.org/10.5281/zenodo.3973172.

## Code availability

NeatMS is open-source and is freely available at https://github.com/bihealth/NeatMS under permissive MIT license. A pypi package is available at https://pypi.org/project/NeatMS/, a Bioconda package is available at https://anaconda.org/bioconda/neatms. The user documentation can be found at https://neatms.readthedocs.io/en/latest/.

## Acknowledgements

The authors thank Alina Eisenberger and Raphaela Fritsche for generating datasets 1 and 2, Friederike Gutmann and Mathias Kuhring for testing NeatMS and providing valuable feedback, Eric Blanc for his valuable insight on machine learning.

## References

1. Smith, C. A., Want, E. J., O’Maille, G., Abagyan, R. & Siuzdak, G. XCMS:□ Processing Mass Spectrometry Data for Metabolite Profiling Using Nonlinear Peak Alignment, Matching, and Identification. Anal. Chem. 78, 779–787 (2006).

2. Pluskal, T., Castillo, S., Villar-Briones, A. & Orešič, M. MZmine 2: Modular framework for processing, visualizing, and analyzing mass spectrometry-based molecular profile data. BMC Bioinformatics 11, 395 (2010).

3. Protsyuk, I. et al. 3D molecular cartography using LC–MS facilitated by Optimus and ’ili software. Nature Protocols 13, 134–154 (2018).

4. Myers, O. D., Sumner, S. J., Li, S., Barnes, S. & Du, X. One Step Forward for Reducing False Positive and False Negative Compound Identifications from Mass Spectrometry Metabolomics Data: New Algorithms for Constructing Extracted Ion Chromatograms and Detecting Chromatographic Peaks. Anal. Chem. 89, 8696–8703 (2017).

5. Myers, O. D., Sumner, S. J., Li, S., Barnes, S. & Du, X. Detailed Investigation and Comparison of the XCMS and MZmine 2 Chromatogram Construction and Chromatographic Peak Detection Methods for Preprocessing Mass Spectrometry Metabolomics Data. Anal. Chem. 89, 8689–8695 (2017).

6. Zhang, A., Lipton, Z. C., Li, M. & Smola, A. J. Dive into Deep Learning. (2020).

7. Borgsmüller, N. et al. WiPP: Workflow for Improved Peak Picking for Gas Chromatography-Mass Spectrometry (GC-MS) Data. Metabolites 9, 171 (2019).

8. Lebanov, L., Tedone, L., Ghiasvand, A. & Paull, B. Random Forests machine learning applied to gas chromatography – Mass spectrometry derived average mass spectrum data sets for classification and characterisation of essential oils. Talanta 208, 120471 (2020).

9. Melnikov, A. D., Tsentalovich, Y. P. & Yanshole, V. V. Deep Learning for the Precise Peak Detection in High-Resolution LC–MS Data. Analytical Chemistry 92, 588–592 (2020).

10. Kantz, E. D., Tiwari, S., Watrous, J. D., Cheng, S. & Jain, M. Deep Neural Networks for Classification of LC-MS Spectral Peaks. Anal. Chem. 91, 12407–12413 (2019).

11. Rong, Z. et al. NormAE: Deep Adversarial Learning Model to Remove Batch Effects in Liquid Chromatography Mass Spectrometry-Based Metabolomics Data. Anal. Chem. 92, 5082–5090 (2020).

12. Rawat, W. & Wang, Z. Deep Convolutional Neural Networks for Image Classification: A Comprehensive Review. Neural Comput 29, 2352–2449 (2017).

13. AbsoluteIDQ^®^ p400 HR Kit - Metabolomics kit for Exactive™. biocrates life sciences ag https://biocrates.com/absoluteidq-p400-hr-kit/ (2020).

14. Tautenhahn, R., Böttcher, C. & Neumann, S. Highly sensitive feature detection for high resolution LC/MS. BMC Bioinformatics 9, 504 (2008).

15. Grüning, B. et al. Bioconda: sustainable and comprehensive software distribution for the life sciences. Nature Methods 15, 475–476 (2018).

16. Chambers, M. C. et al. A cross-platform toolkit for mass spectrometry and proteomics. Nature Biotechnology 30, 918–920 (2012).

17. Perez, F. & Granger, B. E. IPython: A System for Interactive Scientific Computing. Computing in Science Engineering 9, 21–29 (2007).

18. Pan, S. J. & Yang, Q. A Survey on Transfer Learning. IEEE Transactions on Knowledge and Data Engineering 22, 1345–1359 (2010).

19. van der Walt, S., Colbert, S. C. & Varoquaux, G. The NumPy Array: A Structure for Efficient Numerical Computation. Computing in Science Engineering 13, 22–30 (2011).

20. Pedregosa, F. et al. Scikit-learn: Machine Learning in Python. Journal of Machine Learning Research 12, 2825–2830 (2011).

21. Chollet, F. & others. Keras. (2015).

22. Martín Abadi et al. TensorFlow: Large-Scale Machine Learning on Heterogeneous Systems. (2015).

23. McKinney, W. Data Structures for Statistical Computing in Python. in Proceedings of the 9th Python in Science Conference (eds. Walt, S. van der & Millman, J.) 56–61 (2010). doi:10.25080/Majora-92bf1922-00a.

24. Libiseller, G. et al. IPO: a tool for automated optimization of XCMS parameters. BMC Bioinformatics 16, 118 (2015).

