## Supplementary materials for "Deep Learning assisted Peak Curation for large scale LC-MS Metabolomics"

### Supplementary material

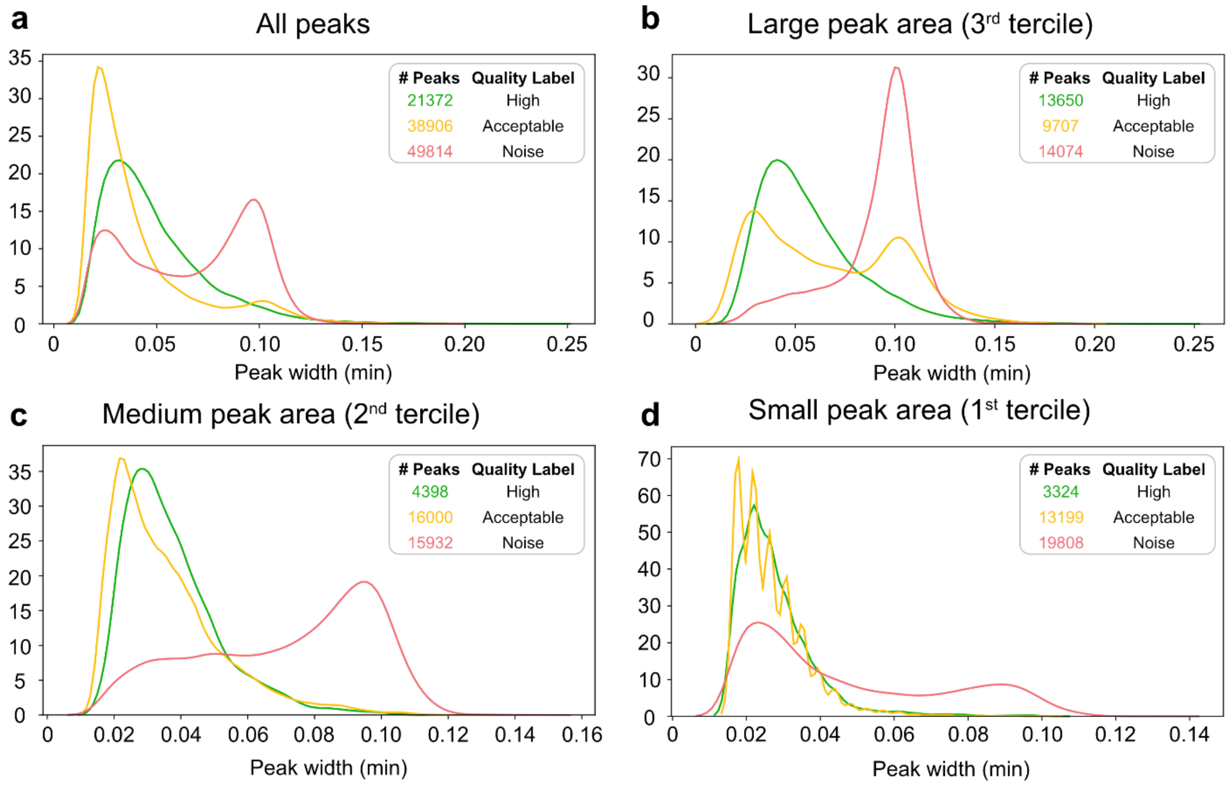


Figure S1: Density plots representing the peak width distribution of the 3 peak classes of dataset 1 analysed using MZmine and NeatMS TL model. a. Width distribution of all peaks. b. Width distribution of peaks from the upper tercile of peak area. c. Width distribution of peaks from the intermediate peak area tercile. d. Width distribution of peaks from the lower peak area tercile.

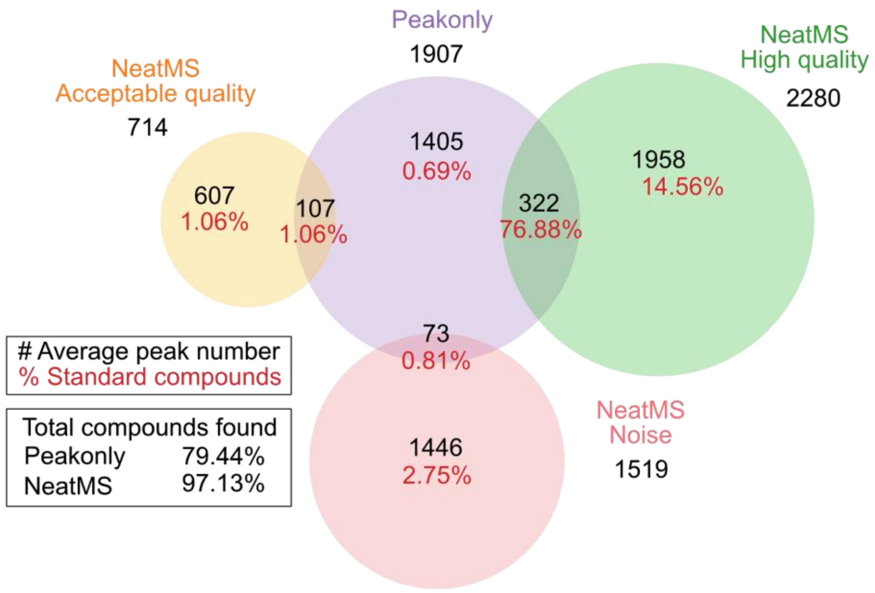


Figure S2: Venn diagram comparing peakonly and the combination XCMS with NeatMS TL model. Numbers are averages over the 20 samples of dataset 1: total number of detected peaks (black), percent of recovered SSCs (red).

a

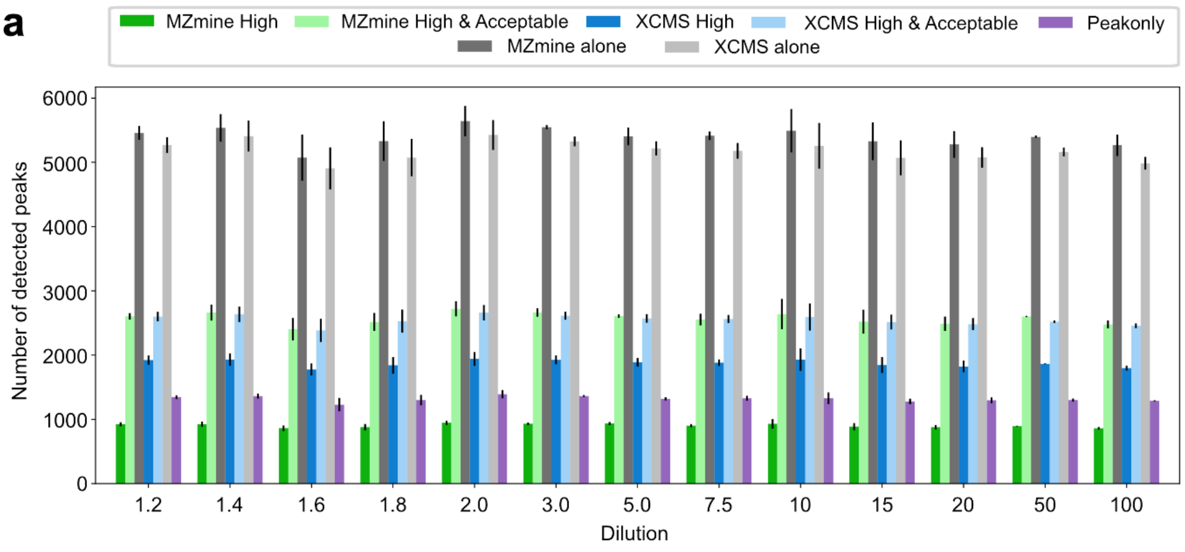


b

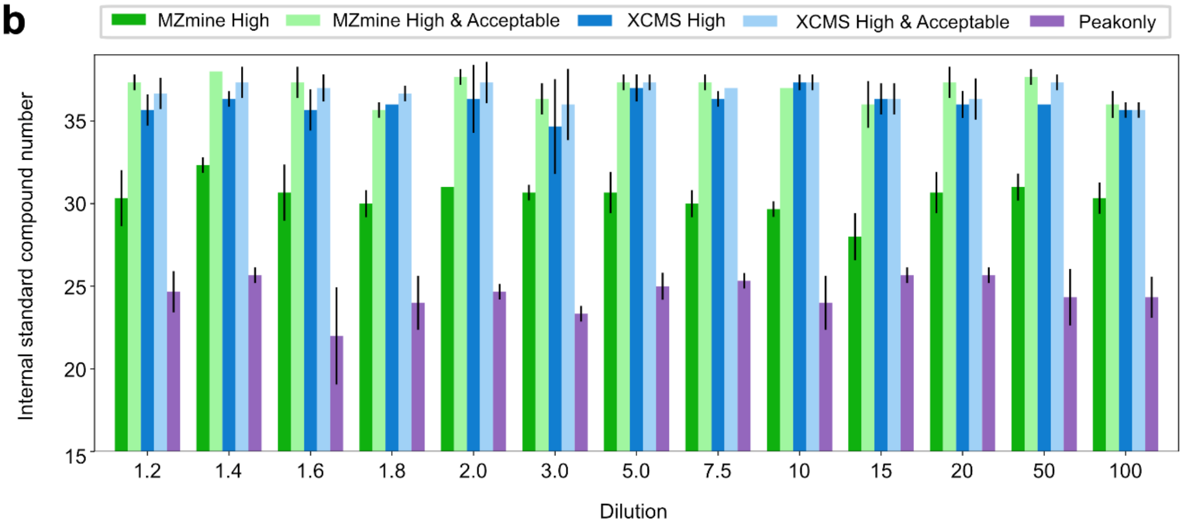


Figure S3: Peak number and internal standard recovery for different tools on the different dilution points of dataset 2. a. Average number of peaks for each dilution point before and after using NeatMS. Only the high quality and acceptable quality classes of NeatMS are displayed here. The difference between the total number of peaks detected by MZmine and XCMS and the number of peaks included in NeatMS high and acceptable quality corresponds to peaks predicted as Noise and peaks rejected by the minimum scan number filter (set to the default value of 5). b. Number of (non diluted) internal standard compounds recovered by the different tools and the details of their predicted classes for each dilution point.

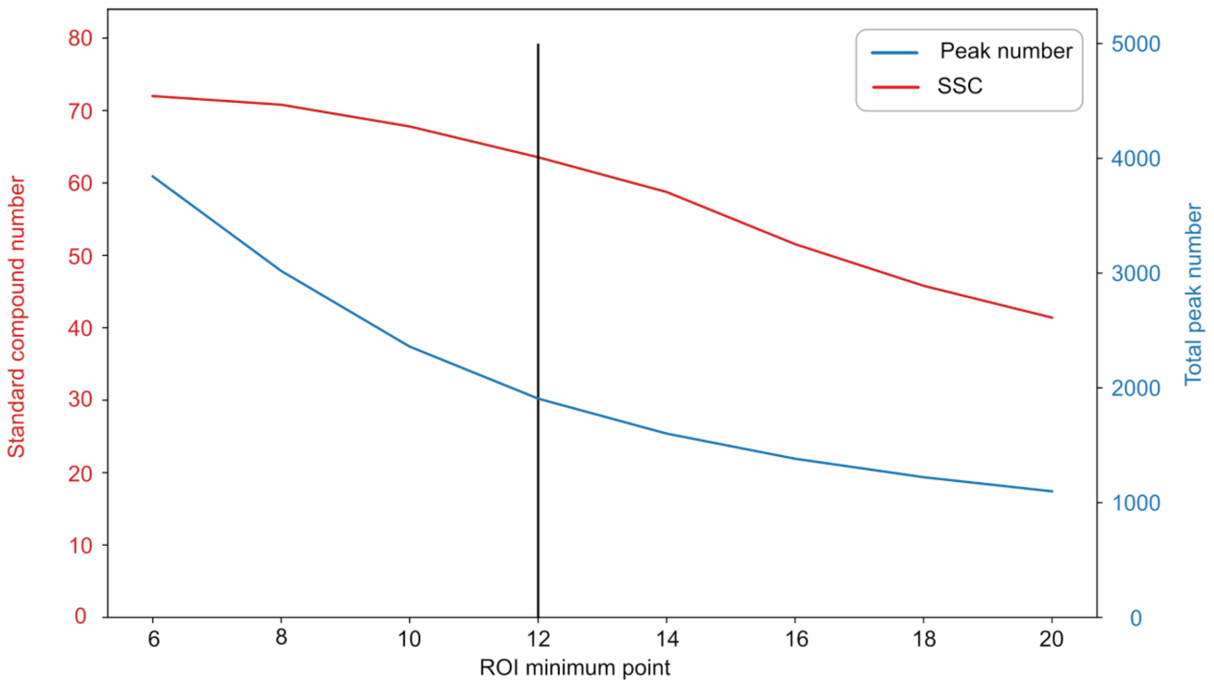


Figure S4: Average detected peak number and standard compound recovered by peakonly on dataset 1 when modulating peakonly “ROI minimum points” parameter. As previously described by the authors of peakonly, we found that lowering the ROI minimum points parameter significantly increases the number of reported noise peaks. A similar effect can be observed with peakonly “Peak minimum points” parameter (not shown). We conclude that the recommendations of peakonly authors are a good compromise between SSC sensitivity and false positive peak detection. The vertical line (in black) represents the parameter that we have used for comparison.

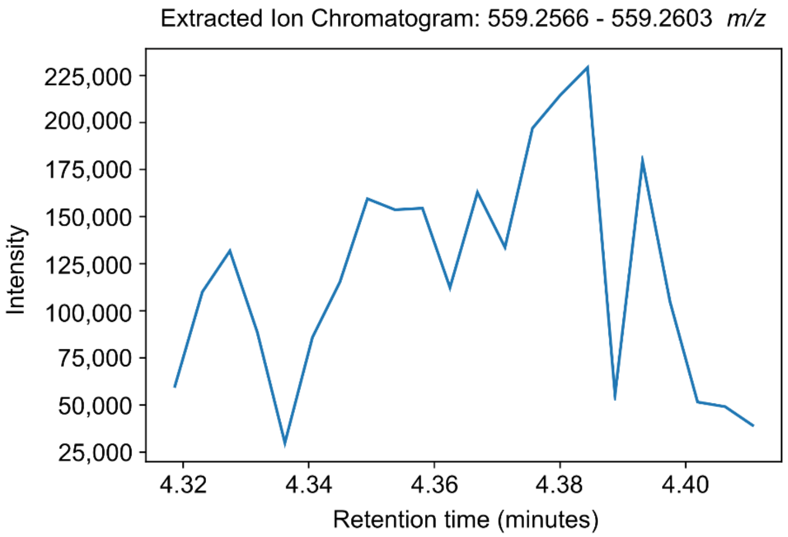


Figure S5: Example of an extracted ion chromatogram of a peak reported by MZmine and classified as Noise by NeatMS matching a CS in dataset 1.

Table S1: Parameters of the different tools used for dataset 1 and 2. Only non default parameters are shown. Peakonly stable and published version 0.1 was used through command lines. XCMS version 3.10.0 was used in R 4.0.1. MZmine was used under version 2.53. NeatMS was used under version 0.6 with default parameters only.

| Parameter | Value |
|---|---|
| MZmine (Dataset 1 & 2) | |
| ADAP chromatogram builder | |
| Min group size in # of scans | 5 |
| Group intensity threshold | $5 \times 10^{2}$ |
| Min highest intensity | $1 \times 10^{3}$ |
| m/z tolerance (m/z) | $1 \times 10^{-2}$ |
| Wavelets (ADAP) chromatogram deconvolution | |
| S/N threshold | 10 |
| S/N estimator | Intensity window SN |
| Min feature height | $1 \times 10^{3}$ |
| Peak duration range | 0.02 - 1.0 |
| RT wavelet range | 0.001 - 0.05 |
| XCMS CentWave (Dataset 1) | |
| Min-Max peak width | 3-85.16 |
| ppm | 17 |
| mzdiff | -0.01145 |
| XCMS CentWave (Dataset 2) | |
| Min-Max peak width | 3-81 |
| ppm | 11.75 |
| mzdiff | -0.016 |
| Peakonly (Dataset 1 & 2) | |
| Delta mz | 0.01 |
| ROI minimum points | 12 |
| Peak minimum points | 6 |

Table S2: Performance of peakonly with NeatMS using dataset 1 and the two models: Average number of peaks across 20 samples, average percentages of the 80 SC recovered using peakonly and different models. The input row shows the results returned by peakonly alone, other rows show the details of the three peak classes given by NeatMS. The total number of peaks after classification is smaller than the input due to the application of a minimum scan number filter that NeatMS uses (default value of 5 is used).

| | | NeatMS peakonly TL model | NeatMS peakonly PT model |
|---|---|---|---|
| Peak number | Input | 1907 | 1907 |
| | Classified | 1724 | 1724 |
| | High Quality | 457 | 730 |
| | Acceptable Quality | 747 | 684 |
| | Noise | 520 | 310 |
| CS found | Input & classified | 79.44% | 79.44% |
| | High Quality | 44.75% | 74.69% |
| | Acceptable Quality | 33.06% | 3.81% |
| | Noise | 1.44% | 0.75% |

#### NeatMS external links to material:

Documentation: https://neatms.readthedocs.io/
Pypi package: https://pypi.org/project/NeatMS/
Bioconda package: https://anaconda.org/bioconda/neatms
Source code: https://github.com/bihealth/NeatMS
Jupyter notebook tutorials: https://github.com/bihealth/NeatMS/tree/master/notebook/tutorial
Test data: https://github.com/bihealth/NeatMS/tree/master/data/test_data
Neural network models: https://github.com/bihealth/NeatMS/tree/master/data/model
Licence: https://github.com/bihealth/NeatMS/blob/master/LICENSE
